# Identification of differentially expressed small RNAs and prediction of target genes in Italian Large White pigs with divergent backfat deposition

**DOI:** 10.1101/245670

**Authors:** Roberta Davoli, Enrico Gaffo, Martina Zappaterra, Stefania Bortoluzzi, Paolo Zambonelli

**Author notes:** Equally contributing first Author (Roberta Davoli, Enrico Gaffo). Equally contributing last Author (Stefania Bortoluzzi, Paolo Zambonelli).

## Abstract

The identification of the molecular mechanisms regulating pathways associated to the potential of fat deposition in pigs can lead to the detection of key genes and markers for the genetic improvement of fat traits. MicroRNAs (miRNAs) interactions with target RNAs regulate gene expression and modulate pathway activation in cells and tissues. In pigs, miRNA discovery is well far from saturation and the knowledge of miRNA expression in backfat tissue and particularly of the impact of miRNA variations are still fragmentary. We characterized by RNAseq the small RNAs (sRNAs) expression profiles in Italian Large White pig backfat tissue. Comparing two groups of pigs divergent for backfat deposition, we detected 31 significant differentially expressed (DE) sRNAs, 14 up-regulated (including ssc-miR-132, ssc-miR-146b, sscmiR-221–5p, ssc-miR-365–5p, and the moRNA ssc-moR-21–5p) and 17 down-regulated (including ssc-miR-136, ssc-miR-195, ssc-miR-199a-5p, and ssc-miR-335). To understand the biological impact of the observed miRNA expression variations, we used the expression correlation of DE miRNA target transcripts expressed in the same samples to define a regulatory network of 193 interactions between DE miRNAs and 40 DE target transcripts showing opposite expression profiles and being involved in specific pathways. Several miRNAs and mRNAs in the network resulted to be expressed from backfat related pig QTLs. These results are informative on the complex mechanisms influencing fat traits, shed light on a new aspect of the genetic regulation of fat deposition in pigs, and facilitate the perspective implementation of innovative strategies of pig genetic improvement based on genomic markers.

## Introduction

Backfat deposition in pigs is an important selection trait closely connected with carcass and meat quality. MicroRNAs (miRNAs) and miRNA-like short RNAs (sRNAs) represent a functionally relevant RNA category modulating the expression of coding messenger RNAs (mRNAs) and noncoding transcripts in almost all tissues. In pigs, only 411 mature miRNAs are included in the current miRBase release 21 (Kozomara and Griffiths-Jones, 2013), being less than 16% of the number of human miRNAs, indicating that miRNA discovery in pigs is far from reaching saturation. To date, only few papers describing miRNAs expression in porcine adipose tissues have been published (Li et al., 2011; Chen et al, 2012; Li et al., 2012; Bai et al., 2014). Recently, the transcription profiles of sRNAs in backfat of two Italian Large White pigs have been described in Gaffo et al. (2014), reporting new miRNAs, isomiRs and moRNAs, and showing the complexity of sRNA population in the porcine backfat tissue. RNA-seq was recently used to study transcriptome variations in backfat tissue comparing two groups of Italian Large White pigs divergent for backfat deposition, getting a deeper knowledge of the molecular processes regulating the potential of fat deposition in pigs (Zambonelli et al., 2016), that is essential to identify key genes and markers for the genetic improvement of fat traits.

In this study, we used RNA-seq to describe the transcription profile of sRNAs expressed in porcine backfat tissue and to identify sRNAs differentially expressed between fat and lean pigs. Furthermore, linking miRNAs and target transcripts whose expression vary in relation to backfat deposition we reconstructed a regulatory network that disclose the impact of miRNAs regulatory action on fat traits.

## Materials and Methods

### Samples collection and sequencing

For this analysis we used 18 of the Italian Large White pigs and the backfat samples collected and described in Zambonelli et al. (2016). All animals used in this study were kept according to Italian and European laws for pig production and all the adopted procedures were fully compliant with national and European Union regulations for animal care and slaughtering. We considered short RNA-seq data of backfat tissue in 18 animals, comprising 9 pigs with high backfat thickness (FAT) and 9 pigs with low backfat thickness (LEAN), as described before in Zambonelli et al. (2016). The 18 RNA samples were newly sequenced (series record GSE108829) as detailed below.

Backfat samples stored at –80°C in a deep freezer were used for total RNA extraction using Trizol (Invitrogen) according to the manufacturer’s instructions. Extracted RNA was quantified using the Nanodrop ND-1000 spectrophotometer, and the quality of the extracted RNA was assayed using an Agilent 2100 BioAnalyzer (Supplementary Table 1). Eighteen small RNA libraries were prepared from total RNA using the TruSeq Small RNA kit (Illumina) and version 3 of the reagents following the manufacturer’s suggested protocol. The libraries were individually tagged and run on 9 lanes of an Illumina GAII.

### Computational analysis of small RNA sequencing data

The bioinformatics pipeline used is summarized in Supplementary Figure 1, informing also on the study design. RNA-seq data were processed with a combination of the miR&moRe pipeline (Bortoluzzi et al., 2012; Gaffo et al., 2014) and of miRDeep2 (Friedländer et al., 2011), which was exploited to discover new precursor hairpins that express mature miRNAs (NPmiRNAs) in the samples: we run miR&moRe on the known porcine miRNAs plus a high confidence set of predictions, obtained by miRDeep2.

**Figure 1.**
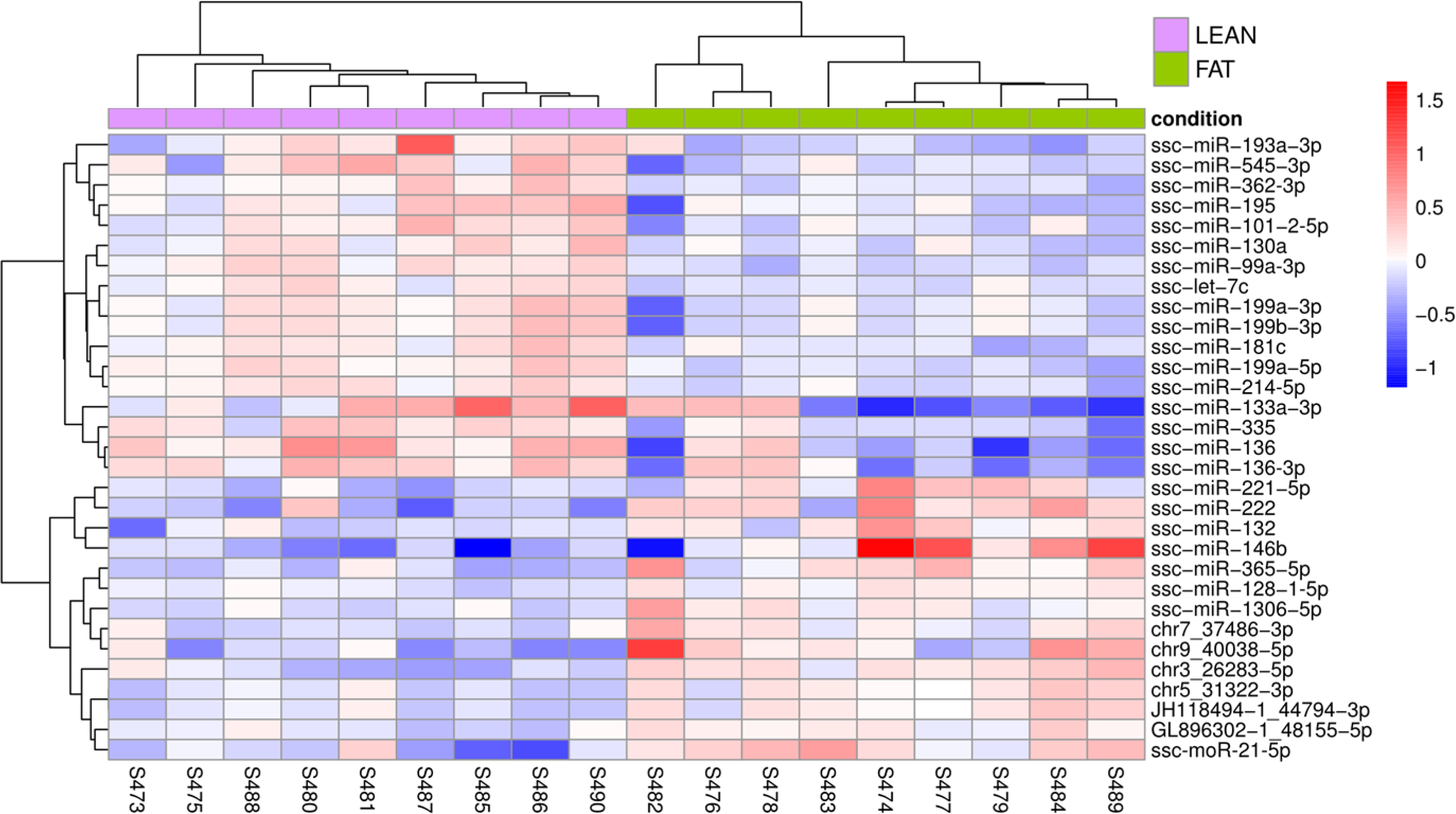
Heatmap of expression profiles of miRNAs differentially expressed (DEMs) comparing FAT vs LEAN animals. Rows of the heatmap represent the 31 DEMs, columns correspond to samples. The heatmap cells are colored according to the deviance of the sRNA expression in the sample from the average expression of the sRNA, thus red and blue cells represent respectively expression values higher or lower than mean expression across all samples (white) with color intensity proportional to the difference from the mean, in the regularized logarithmic scale. **File: Figure1_rev.pdf**

MiRDeep2 v0.0.5 was run with default parameters pooling the sample read sets. Input genome and miRNA annotation were the same used for miR&moRe (UCSC database susScr3 genome and miRBase release 21). We filtered miRDeep2 predictions restricting to precursors with miRDeep2 score greater than 1.0, length greater than or equal to 50 nt and predicted probability of being a miRNA greater than 60% (Londin et al., 2015). The novel precursor annotation predicted by miRDeep2 was added to miRBase swine miRNA annotation and given as input to miR&moRe, which was run on the 18 newly sequenced samples.

MiR&moRe quantifies small RNAs from RNA-seq experiments, detect as well miRNA isoforms (isomiRs), novel miRNAs expressed in known hairpin precursors with only one miRNA annotated, and identify also miRNA-offset RNAs (moRNAs; Bortoluzzi et al., 2011; Bortoluzzi et al., 2012; Guglielmelli et al., 2015). The miR&moRe pipeline performs a preliminary cleaning and quality preprocessing of the input raw sequences. Reads passing the quality filter are aligned to the reference miRNA precursors and to the reference genome for expression quantification. Identification and expression quantification of isomiRs and moRNAs follow from the alignments and sequence folding predictions. Small RNAs expression levels are measured as read alignment counts in each sample.

### Differential expression assessment and validation

Read counts were normalized across all the samples according to the DESeq2 (v1.4.5; Love et al., 2014) approach. Small RNAs represented by less than ten normalized reads considering all samples were excluded from further analysis. Short RNA differential expression comparing FAT and LEAN groups was assessed by DESeq2, considering significant those variations with FDR <0.05 (Benjamini-Hochberg adjusted P-values).

Validations of miRNAs differentially expressed according to RNA-seq analysis have been obtained by Quantitative Real Time PCR (qRT-PCR) analysis on the same samples, performed in triplicate. The TaqMan^®^ Micro RNA Assay kit (Applied Biosystems) was used, following the customer protocol, as reported in Gaffo et al. (2014), and using as internal reference the U6 snRNA (Chen et al., 2012; Yu et al., 2012). For the relative miRNA quantification, the PCR-derived cycle threshold (Cq) of target miRNAs is compared with that of a stably expressed endogenous miRNA from the same sample. The difference between these values is the ΔCq value (Rai et al., 2012). A t-test was used to assess the significance of the differential expression estimated by qRT-PCR, considering a significance threshold of P-value <0.05.

### Small RNA target prediction on backfat long RNA data

The transcription profiles of long RNAs in the same pig backfat samples (GSE68007) already analyzed in Zambonelli et al. (2016) were used for integrative analyses with miRNA expression profiles, considering matched miRNA and transcript expression data in backfat of FAT and LEAN animals. MiRanda v3.3a (Enright et al., 2003) was applied to predict target sites on the 63,418 transcript sequences reported by Zambonelli et al. (2016). For each differentially expressed miRNA (DEM), the prediction was computed using the most abundant isomiR sequences (detected as reported in Gaffo et al., 2014), keeping only isomiRs contributing each at least 10% of the miRNA expression.

Next, focusing on differentially expressed small RNAs, we calculated Spearman pairwise correlations between sRNA and DEM expression profiles, and tested the significance of the correlations. According to largely prevalent repressive role of miRNAs on target transcripts, we focused on the negative correlations, whose association P-values were corrected for multiple tests (Benjamini-Hochberg). Only negative correlations with FDR at most 10% were considered significant to identify miRNA-transcript relations that are supported by expression data analysis. Significant miRNA-transcript relations with correlation <-0.6 were selected to draw an interaction network, using Cytoscape version 3.3. Pathways enriched in genes included in the network have been calculated with EnrichR (http://amp.pharm.mssm.edu/Enrichr/; Kuleshov et al., 2016), selecting KEGG and Reactome databases and considering significant those pathways with adjusted P-value at most 0.1 and including at least two genes.

### Quantitative Trait Loci enrichment on target genomic regions

For each pig QTL annotated in the PigQTL database (http://www.animalgenome.org/cgi-bin/QTLdb/SS/download?file=gbpSS_10.2), we counted the number of differentially expressed sRNAs and their predicted target genes considering only those with differentially expressed transcripts with FDR at most 30% according to Zambonelli et al. (2016) mapping in the genomic region. QTL enrichment in miRNA and genes was tested using the upper-tailed hypergeometric test for over-representation, with one side mid p-values as defined in Rivals et al. (2007). The analysis used as background the pig genome annotation (v 10.2.80) merged with the new genes discovered in Zambonelli et al. (2016), resulting in 34,617 genes, and the new sRNAs detected in this study. We considered as significantly enriched only those QTLs showing adjusted P-values <0.1.

## Results

### Small RNA identification and quantification from RNA-seq data

RNA-seq analysis produced in 97 million reads per sample on average, that after read trimming and subsequent filtering steps (Supplementary Table 2) were reduced to 37 million high quality reads per sample, in average, that were further processed for the sRNAs characterization. We applied stringent filtering criteria in the processing of the raw data to obtain high quality read sets deprived of sequencing artifacts, at the cost of high number of discarded sequences. This approach and the number of biological replicates contributed to the reliability of downstream predictions, in particular regarding novel elements like moRNAs and isomiRs.

MiRDeep2 analysis of RNA-seq data predicted 1,340 new miRNA precursors characteristic of pig genome, expressing 2,680 putative novel mature miRNAs, whose sequences do not overlap each other and with different sequence from pig miRNA precursors reported in miRBase. Of these, 103 NPmiRNAs passed the filtering criteria and were considered for following analyses. Overall, 426 sRNAs resulted expressed in the new dataset, including 231 known miRNAs, 69 new miRNAs from known precursors, 103 new miRNAs from new precursors, and 23 moRNAs (Supplementary Table 3).

As observed in Gaffo et al. (2014), the expression distribution was very skewed: only 24 known miRNAs (5.6% of the expressed sRNAs) accounted for the 90% of the total expression, and only 21.1% of expressed sRNAs accounted the 99% of the total expression (Supplementary Figure 2). Ssc-miR-10b, ssc-miR-143–3p and ssc-148a-3p resulted very abundant and accounted together for about the 52% of the total expression, with the first two accounting for respectively about 33% and 13% of the expression.

### Differentially expressed sRNAs

Among the 426 sRNAs expressed in the considered porcine backfat samples, 31 resulted differentially expressed comparing LEAN and FAT animals (Table 1, Supplementary Table 4): 18 known miRNAs, 6 new sister miRNAs, 6 miRNAs from new precursors and ssc-moR-21–5p, whose existence was previously validated by qRT-PCR (Gaffo et al., 2014). Figure 1 shows the unsupervised clustering and heatmap of expression profiles of the 31 DEM in the 18 samples and two animal groups considered. SRNAs up- and down-regulated in FAT animals were 14 and 17, respectively. Up-regulated sRNAs include six known miRNAs (ssc-miR-146b, ssc-miR-365–5p, ssc-miR-221–5p, ssc-miR-222, ssc-miR-132 and ssc-miR-1306–5p) the moRNA ssc-moR-21–5p, a new pig sister miRNA (ssc-miR-128–1-5p) and six mature miRNAs from newly predicted precursors. All down-regulated sRNAs are miRNAs from already annotated precursors, including 12 known miRNAs (ssc-let-7c, ssc-miR-130a, ssc-miR-181c, ssc-miR-199a-5p, ssc-miR-199a-3p, ssc-miR-199b-3p, ssc-miR-195, ssc-miR-193a-3p, ssc-miR-335, ssc-miR-133a-3p, ssc-miR-545–3p, ssc-miR-136) and 5 new sister miRNAs (ssc-miR-214–5p, ssc-miR-99a-3p, ssc-miR-101–2-5p, sscmiR-136–3p, ssc-miR-362–3p, ssc-miR-128–1-5p).

**Table 1.**
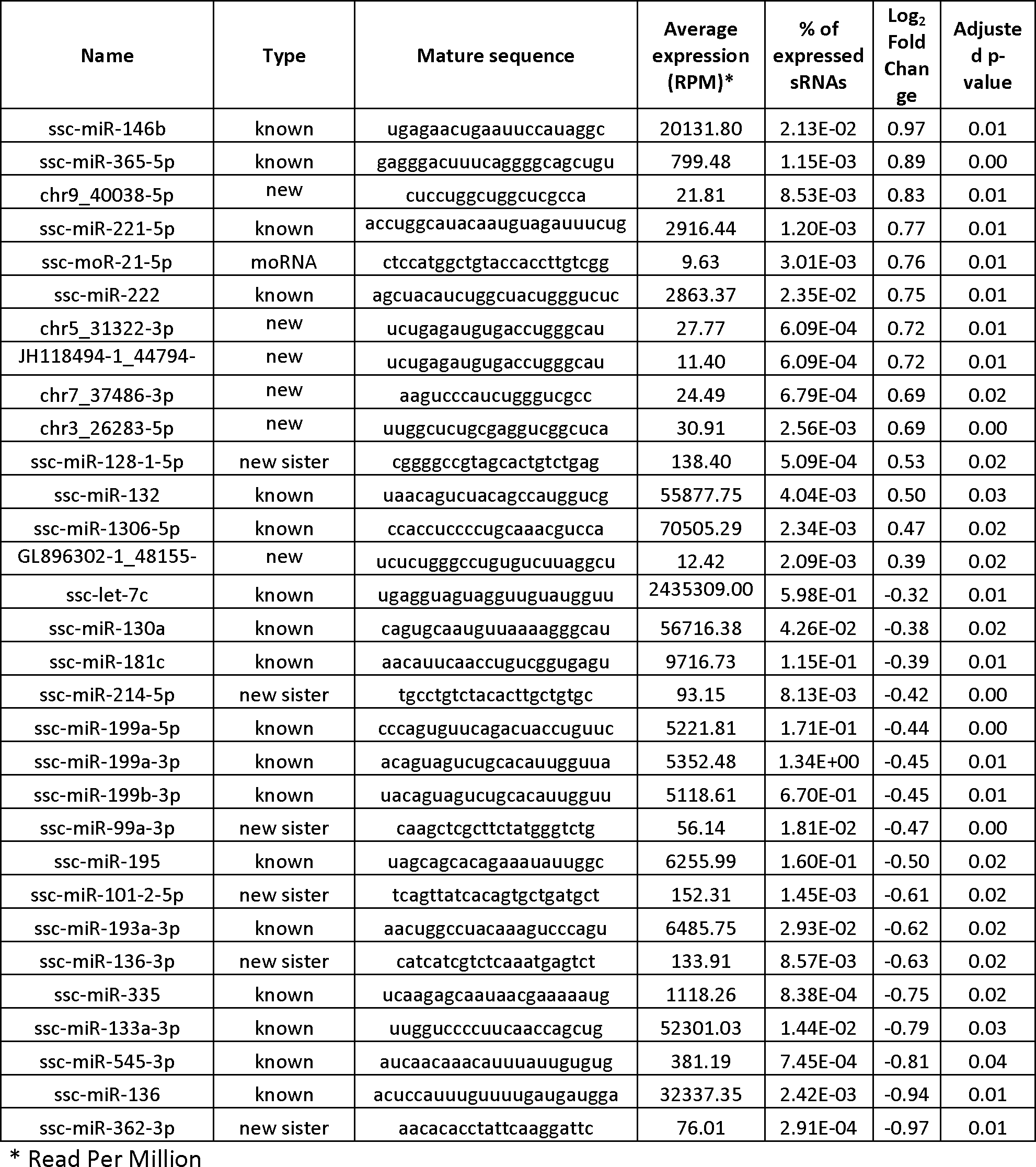
Short RNAs resulting differentially expressed in backfat of FAT vs LEAN animals according to RNA-seq data.

We selected 9 DEMs, 8 miRNAs and ssc-moR-21–5p, for the qRT-PCR validation of differential expression. A good agreement between RNA-seq and qRT-PCR expression estimates was observed (R^2^=0.78, Figure 2A) supporting the robustness of the data reported for all the small RNAs detected in this study. Indeed, all the comparisons performed show the same trend between the two analyses, and 5 out of 9 DE sRNAs resulted significantly differentially expressed also according to RT-PCR estimations (Figure 2B).

**Figure 2.**
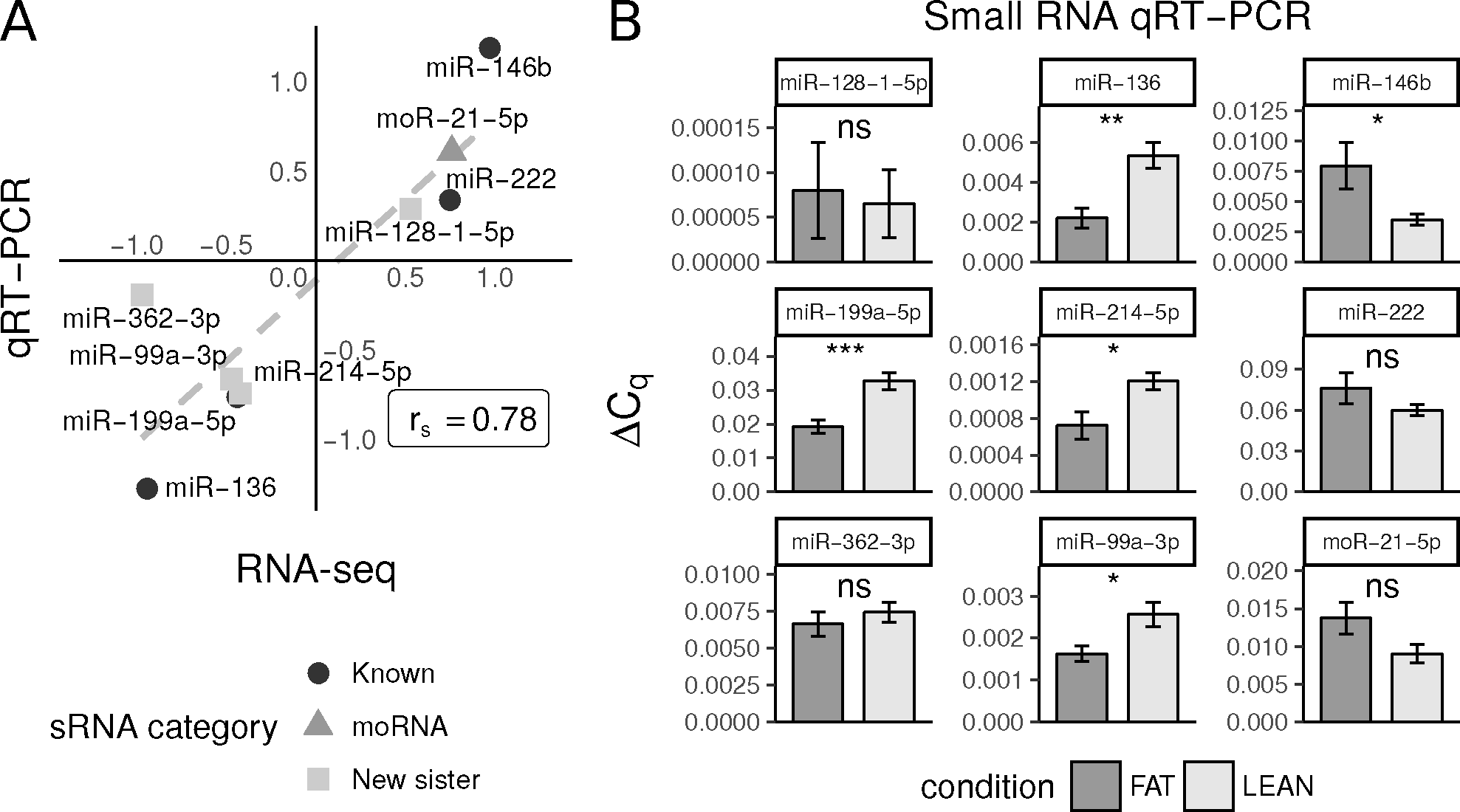
Validation by qRT-PCR of eight DE miRNAs and of the DE moRNA detected by RNA-seq. A) Correlation between fold changes calculated according to RNA-seq (horizontal axis) and qRTPCR (vertical axis) estimations. Dashed line indicates linear smoothing of the points. R_s_ = Spearman’s rank correlation coefficient. B) Significance of differential expression tests according to qRT-PCR data. T-test p-value: ns = not significant; * = <0.05, ** <0.01; *** = <0.001. ΔCq = difference between U6 and the target miRNA PCR-derived cycle thresholds. **File: Figure2_rev.pdf**

### IsomiR composition of differentially expressed sRNAs

IsomiRs were investigated for the DEM showing more than one variant (18 known and 6 novel-precursor miRNAs out of the 31 DEM). Fifty-nine distinct isomiRs were detected, specifically 58 in LEAN and 55 in FAT of which 54 were in common. Four isomiRs are specific for LEAN samples and one is specific for FAT samples (Supplementary Table 5).

The number of isomiRs contributing at least 10% of each DEM expression ranged from one to four, a maximum value detected in three cases (chr5_31322–3p, JH118494–1_44794–3p, chr9_40038–5p). Three miRNAs (chr7_37486–3p, ssc-miR-136 and ssc-miR-193a-3p) are represented by only one major isoform accounting for more than half of the miRNA expression (Supplementary Figure 3A). Most isoforms, according to miR&moRe classification, are variations of the length: isomiRs in the “shorter_or_longer” category are 32 out of 58 in LEAN and 29 out of 55 in FAT; the variants perfectly matching the reference sequence (“exact” variants, or “canonical” isomiRs) are 18 in both groups; and “one-mismatch” variation isomiRs are eight. Less than half of DEMs (10 cases in LEAN and 11 in FAT) express the canonical isomiR as the major form. Conversely, “shorter or longer” isoforms compose the largest part of the expression in 12 LEAN and 11 FAT cases. Notably, in 6 DEMs the exact isomiR contributed less than 10% to the miRNA expression (Supplementary Figure 3B). These findings are consistent with our previous results (Gaffo et al., 2014) showing a similar distribution of isomiRs and indicating that in pig backfat the canonical miRNA isoform is not always the most expressed for the miRNA.

### Network of short and long differentially expressed RNA interactions in pig backfat

The target prediction considered 66 sRNA sequences, including 59 abundant isomiRs from the 18 known miRNAs and the 6 NPmiRNAs, plus 6 new sister-miRNAs and ssc-moR-21–5p. The predicted relations with significant strong negative correlations accounted for 56,683 sRNA-transcript interactions, involving 22,362 transcripts from 12,373 unique genes. All the 66 isomiRs potentially targeted at least one transcript each. Among the known miRNAs, ssc-miR-365–5p presented the largest number of targets (3,095 different transcripts targeted, corresponding to 2,624 genes), while in absolute chr3_26283–5p has the largest number of target transcripts (3,383; 2,878 genes). Ssc-miR-136–3p has the smallest number of targets (524; 465 genes), while among the new miRNAs chr7_37486–3p has the minimum (1,011; 862 genes). Considering the very large number of predicted interactions we focused particularly on the predicted sRNA targets included among the 86 DETs reported in Zambonelli et al. (2016), resulting in 193 putative regulatory interactions between 30 DEMs and 40 target DETs (Supplementary Table 6) that may contribute to regulate the metabolic activity of porcine backfat. The network in Figure 3A-B show the interactions supported by strong correlation. KEGG and Reactome pathways significantly enriched in target DETs in the network are shown in Figure 3C. The down regulated genes *HSPA1A*, *HSPA1B*, and *DNAJB1* are linked to cellular stress responses, to the regulation of stress-induced transcription by *HSF1*, and its modulation by attenuation that occurs during continuous exposure to intermediate heat shock conditions or upon recovery from stress. The same three genes and *MRC1* and *ATP6V0D2* participate in several pathways linked to immunity. Furthermore, five genes (*ADSSL1, AKAP5, BCAT1, HMOX1*, and *PLIN2*) are linked to metabolic pathways.

**Figure 3.**
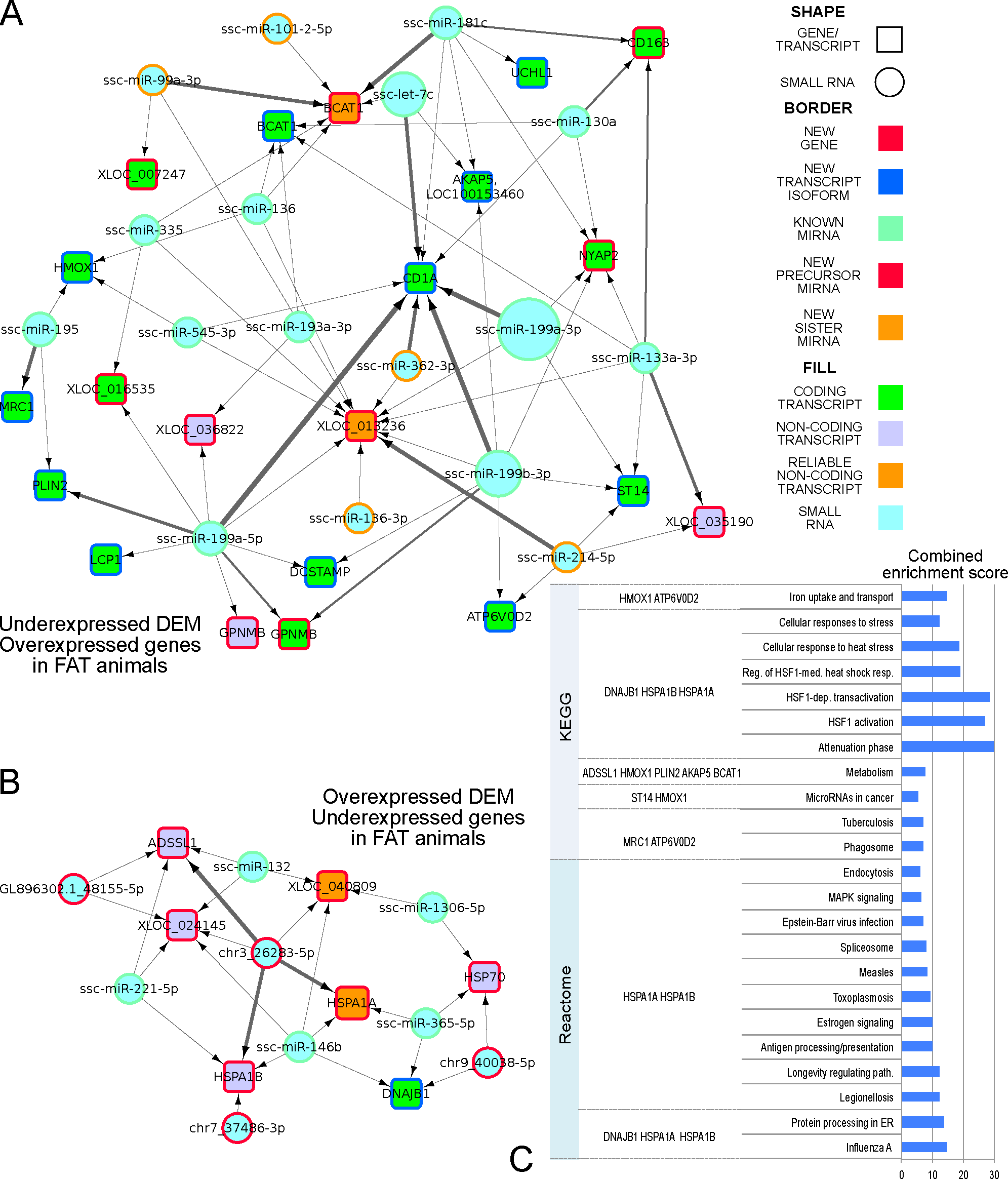
Differentially expressed miRNAs and their putative predicted target transcripts differentially expressed. The network depicts the predicted interactions between. A) miRNAs downregulated in FAT animals and putative target transcripts upregulated in FAT animals; B) miRNAs upregulated in FAT animals and putative target transcripts downregulated in FAT animals; miRNA-target relations with expression profiles correlation <–0.6 are reported, involving 26 miRNA (blue circle nodes) and 28 transcripts (square nodes, coloured green if coding, lilac if non-coding, and orange if reliably non-coding) for a total of 92 interactions (edges). Edge thickness is inversely proportional to the correlation intensity (the smaller the correlation, the thicker the line). Nodes borders are coloured in red if representing a newly predicted transcript or miRNA precursor, orange for new sister miRNAs, and green for known miRNAs. C) KEGG and Reactome pathways significantly enriched in genes included in the network of panel A-B. **File: Figure3_rev.pdf**

The involvement of several genes in the network in specific pathways is informative on the putative regulatory activity of selected miRNAs beyond single targets, suggesting their impact on pathway regulation by coordinated activity on several functionally connected genes. Four of the metabolism-linked genes are overexpressed in FAT animals and resulted under putative control of common underexpressed miRNAs, with both *BCAT1* and *AKAP5* possibly repressed by the ssc-let-7c, and *HMOX1* and *BCAT1* by ssc-miR-195. The underexpressed stress responses genes are under putative control of specific miRNAs, as *HSPA1A*, *HSPA1B*, and *DNAJB1* can be coordinately repressed by the upregulated and validated ssc-miR-146, and the triplet *HSPA1B, DNAJB1* and *HSP70* by ssc-miR-365.

### QTL enrichment of differentially expressed sRNAs and mRNAs loci

As miRNAs and genes with transcripts differentially expressed in LEAN and FAT animals and their interactions could be implicated in genetic modulation of backfat deposition, we looked for the overlap between their loci and pig QTL, and we observed 56 significantly enriched QTL (Supplementary Table 7). Notably, a significant enrichment in miRNAs and their target DE genes was observed for four backfat-specific carcass quality QTLs (Supplementary Table 7). Two largely overlapping QTLs (QTL 21222 “Average backfat thickness” chr4:74,725,606–82,527,275; QTL 21228 “Backfat at first rib” chr4: 76,741,953–83,693,913) covering approximately 9 Mb contains four DE transcripts from *PENK* gene and from a predicted gene. Two enriched “Average backfat thickness” QTLs were identified in chromosome 8 (QTL 29568, chr8: 33,033,004–33,985,796, XLOC_032101 / *UCHL1*) and 12 (QTL 5990, chr12: 23,672,762–60,040,247). The most interesting enriched QTL is located in chromosome 12 and contains the genes expressing eight DETs, and notably five DE miRNAs (the up-regulated ssc-miR-193a-3p and ssc-miR-195 and the down-regulated ssc-miR-132, ssc-miR-365 and ssc-moR-21–5p) strengthening the indication of a possible role of these sRNAs in the backfat phenotype variation.

## Discussion

We reported the landscape of sRNA expression in swine backfat, detecting and quantifying 426 sRNAs expressed, including 403 miRNAs and 23 moRNAs, confirming previous data (Gaffo et al., 2014) and extending them, by increasing the number of sRNAs detected in pig backfat.

The FAT and LEAN animals expression profile comparison disclosed 31 differentially expressed sRNAs whose abundance is related to backfat level, including 30 miRNAs and a moRNA, sscmoR-21–5p, that resulted downregulated in FAT animals. These results are backed up by quantitative real-time PCR validations conducted on a sizeable subset of the DE sRNAs detected by RNA-seq (Figure 2), including known miRNAs, new mature miRNAs of known precursors, and a moRNA, with different expression levels and variable variations strengths in the comparison between FAT and LEAN animals.

Thirthy DE miRNAs possibly targeted 40 out of 86 backfat DETs, while no putative targets were detected among the same backfat DETs for ssc-moR-21–5p. This study confirmed a possible link between ssc-moR-21 and adipocyte physiology (Gaffo et al., 2014) that deserves further investigation.

Altogether, almost two thousands putative miRNA-target interactions emerged considering the correlation between the DE miRNAs and the DE mRNAs and represented as a functional network (Figure 3).

The majority of the transcripts included in the network are coding (green squares) while there are four putative noncoding (grey and orange squares). *BCAT1 (*branched chain amino acid transaminase 1) and *GPNMB (*glycoprotein Nmb) are represented by both coding and one noncoding isoforms.

The less expressed ssc-miR-199a-5p potentially regulate several transcripts, including coding transcripts of different genes, different isoforms derived from the same gene, and unannotated transcripts found overexpressed in FAT vs LEAN pigs. As already described in Zambonelli et al. (2016) some of them are involved in lipid metabolism (perilipin 2, *PLIN2*) or in the regulation of food intake (*GPNMB*). Ssc-miR-199a-5p was found under-expressed in the FAT pigs, in agreement with previous data on the underexpression of this miRNA in attenuating cell proliferation and promoting lipid deposition in porcine pre-adipocytes (Shi X-E et al., 2014). Ssc-miR136, ssc-miR-99a-3p, and ssc-miR-214–5p were down-regulated in FAT vs LEAN pigs. In lambs, miR-136 was differentially expressed in subcutaneous adipose tissue compared with perirenal fat (Meale et al., 2014). Guo et al. (2012) linked miR-99a to preadipocyte differentiation both in porcine intramuscular and subcutaneous vascular stem cells. The same miRNA was twofold more expressed in adult subcutaneous adipose tissue than in juvenile adipose tissue in pigs by Li et al. (2011). miR-214 was previously related to adipose-derived stem cells differentiation in rats) and resulted up-regulated in rat stem cells isolated from subcutaneous fat compared with omentum (Hu et al., 2017).

The importance of ssc-miR-335 in the regulation of biological and molecular processes responsible of different fat deposition between FAT and LEAN animals is supported by previous data linking this miRNA to the control of fat traits (Nakanishi et al., 2009; Oger et al., 2014; and Zhu et al., 2014). In our network, the down-regulation of miR-335 is linked to *BCAT1* overexpression. Under-expressed putative targets of overexpressed miRNAs include one protein coding gene transcripts (*DNAJB1*) and noncoding transcripts, mostly belonging to heat shock protein genes (*HSP70, HSPA1A, HSPA1B*) that deserve further investigation.

Ssc-miR-146b could control the expression of three genes down-regulated all coding for heat shock proteins (*DNAJB1, HSPA1A, HSPA1B*), whereas ssc-miR-365–5p was linked to *DNAJB1, HSPA1A* and *HSP70*, indicating that the two overexpressed *miRNAs* could contribute to regulation pattern of common targets possibly with a synergistic effect. The action of miR-146b on tissue inflammation was described in human obesity by different Authors (Hulsmans et al., 2012; Shi C et al., 2014) who reported that, despite the obese condition, an attenuation of cytokine signaling reduced the effect of the inflammation on the tissues. In visceral and subcutaneous adipose tissue of over weighted and obese human subjects, miR-146b overexpression promoted preadipocyte differentiation, not impacting proliferation (Ahn et al., 2013; Chen et al., 2014). These evidences suggest a likely explanation of the lower level of expression on backfat tissue in FAT pigs of heat shock protein DETs as response to a chronic inflammation (Zambonelli et al., 2016) providing as well a link with miRNA expression and regulatory activity.

Of miRNAs overexpressed in FAT animals, ssc-miR-132 was upregulated in the hypothalamus of rats fed with a high fat diet (Sangiao-Alvarellos et al., 2014), and in omental fat and plasma of obese humans (Heneghan et al., 2011), whereas the overexpression of miR-221 was associated to an increase in adipocitokines expression in mice (Parra et al., 2010) and with body mass index increase in subcutaneous and abdominal adipose tissue in human Pima Indians (Meerson et al., 2013) where, intriguingly, the increase of body mass index was associated also with the under-expression of miR-199a-3p, fully in accordance with our observations.

Several elements of the network, are expressed by genomic regions harboring QTL associated with backfat thickness, supporting the presence and contributing to the identification of positional and functional candidate genes for fat traits. The SSC12 QTL region arbors both the overexpressed ssc-mir-132 and the under-expressed ssc-mir-195. Within a QTL it is possible to found an epistatic effect of different genetic elements acting on opposite direction in the phenotype determination (Cordell, 2002). However, the action on the phenotype of these two miRNAs could be concordant or even synergistic if they repress the expression of genes whose products have opposite roles in biological processes or signaling pathways.

Collectively, our data and the above discussed literature informed on the involvement of miRNAs in the regulation of genes active in adipose tissue, suggesting that the identified DE miRNAs can modulate several functions related to the adipose tissue development and growth, in particular adipocyte differentiation, adipogenesis, lipid deposition, and obesity in humans. Specific miRNAs might modulate as well heat shock transcript expression, supporting the idea that pigs with an increased potential for fat deposition present some aspects similar to those described in human obesity in which a mild inflammation blocks an excessive harmful effect on a high level of inflammation when a chronic increase of fat tissue as in obesity occurs.

We found new genes, mRNAs and miRNAs that, according to sequence and expression data, could be involved in the complex network of regulation of backfat deposition. Our results explain part of the complex mechanisms influencing fat traits, shedding light on new aspect of the genetic regulation of fat deposition in pigs, and facilitate the perspective implementation of innovative strategies of pig genetic improvement based on genomic markers.

## Acknowledgements

This work was supported by Progetto “AGER – Agroalimentare e ricerca”: Advanced research in genomics and processing technologies for the Italian heavy pig production – Hepiget (Grant N. 2011–0279).

The authors declare that they have no competing interests.

## Supporting information

Additional supporting information may be found in the online version of this article.

---

**Supplementary Tables are included in the file: DEM-DET-**

**Target_SupplementaryTables_rev.xlsx**

**Table S1 (ST1).** Samples RIN numbers.

**Table S2 (ST2).** Sample raw reads filtering.

**Table S3 (ST3).** Small RNAs expressed.

**Table S4 (ST4).** List of all DE tests obtained.

**Table S5 (ST5).** IsomiRs specific for FAT or LEAN samples.

**Table S6 (ST6).** Differentially expressed small RNA targets among differentially expressed transcripts.

**Table S7 (ST7).** QTL enrichment for differentially expressed sRNAs and target genes.

**Figure S1.**
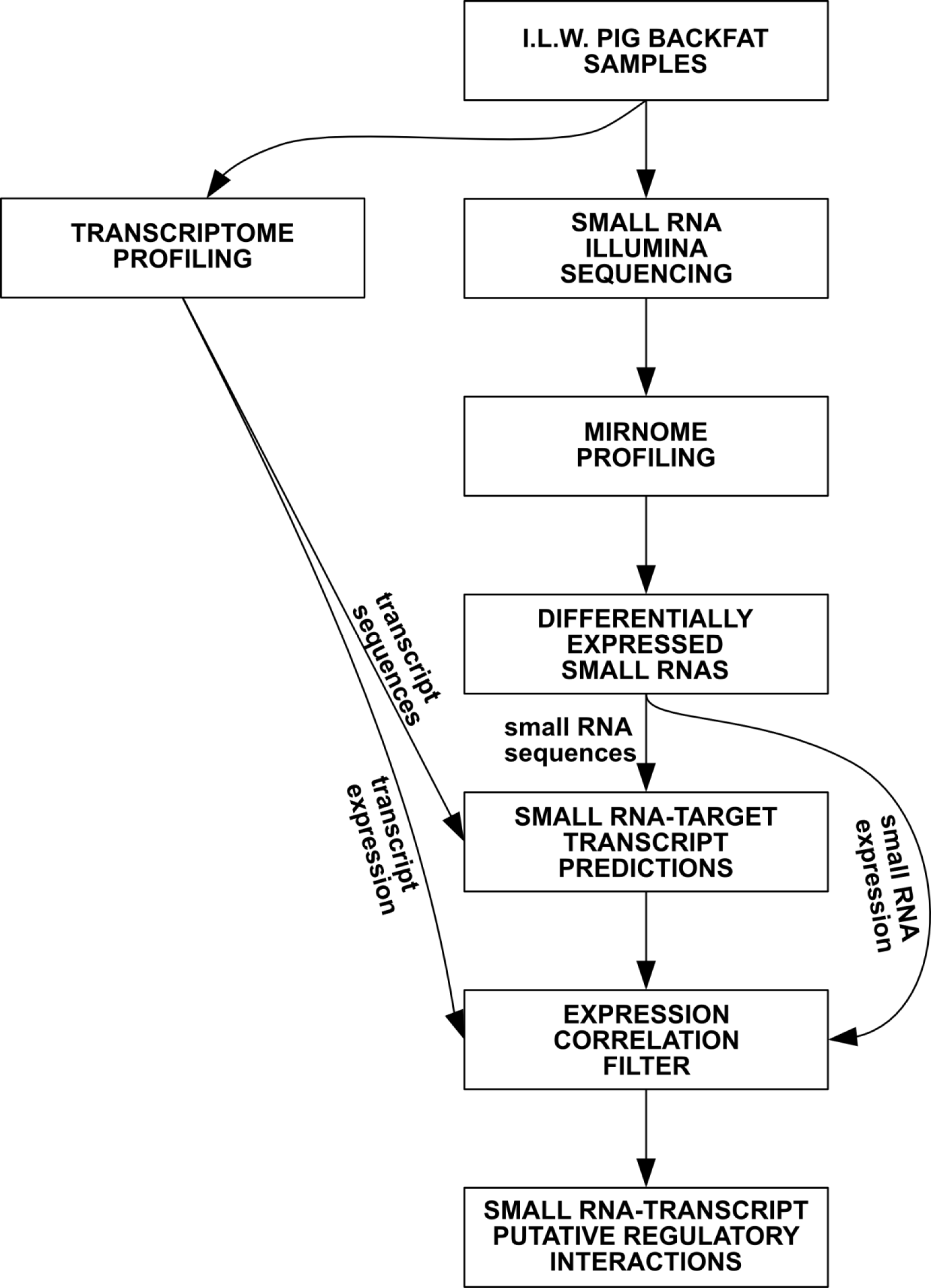
Outline of the computational analysis workflow. Long and short RNA-seq data in the same backfat samples of 10 LEAN and 10 FAT animals have been considered; after detection, discovery and quantification of miRNA and miRNA-like small RNAs, integrative analyses of target prediction and of miRNA and transcript expression profiles were used for miRNA-transcript regulatory network inference. File: FigureSF1_rev.pdf

**Figure S2.**
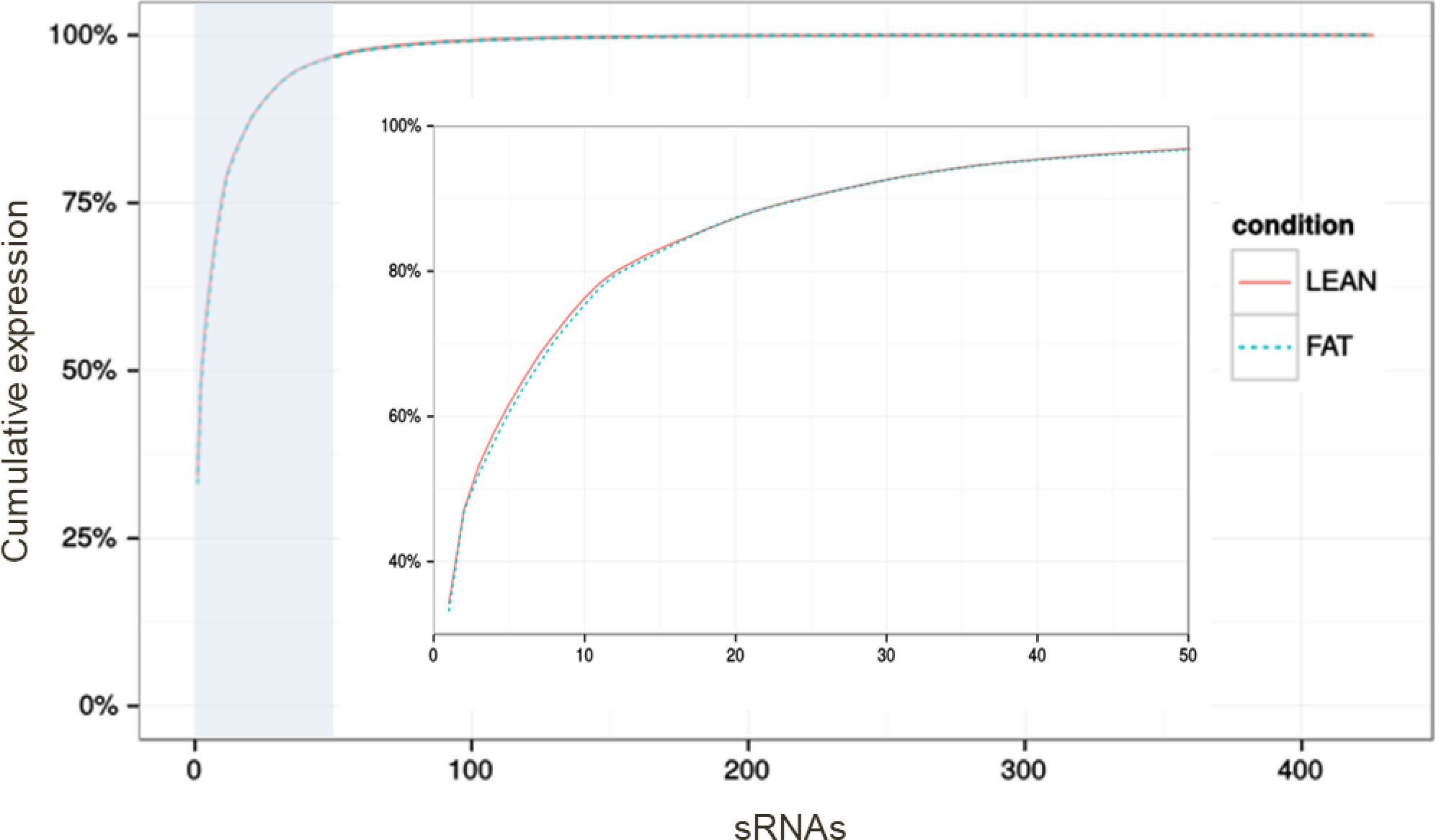
Cumulative expression of gene expression measures show a similar pattern in the FAT and LEAN groups. The small panel inside the figure corresponds to the region shaded in light blue in the main panel. File: FigureSF2_rev.pdf

**Figure S3.**
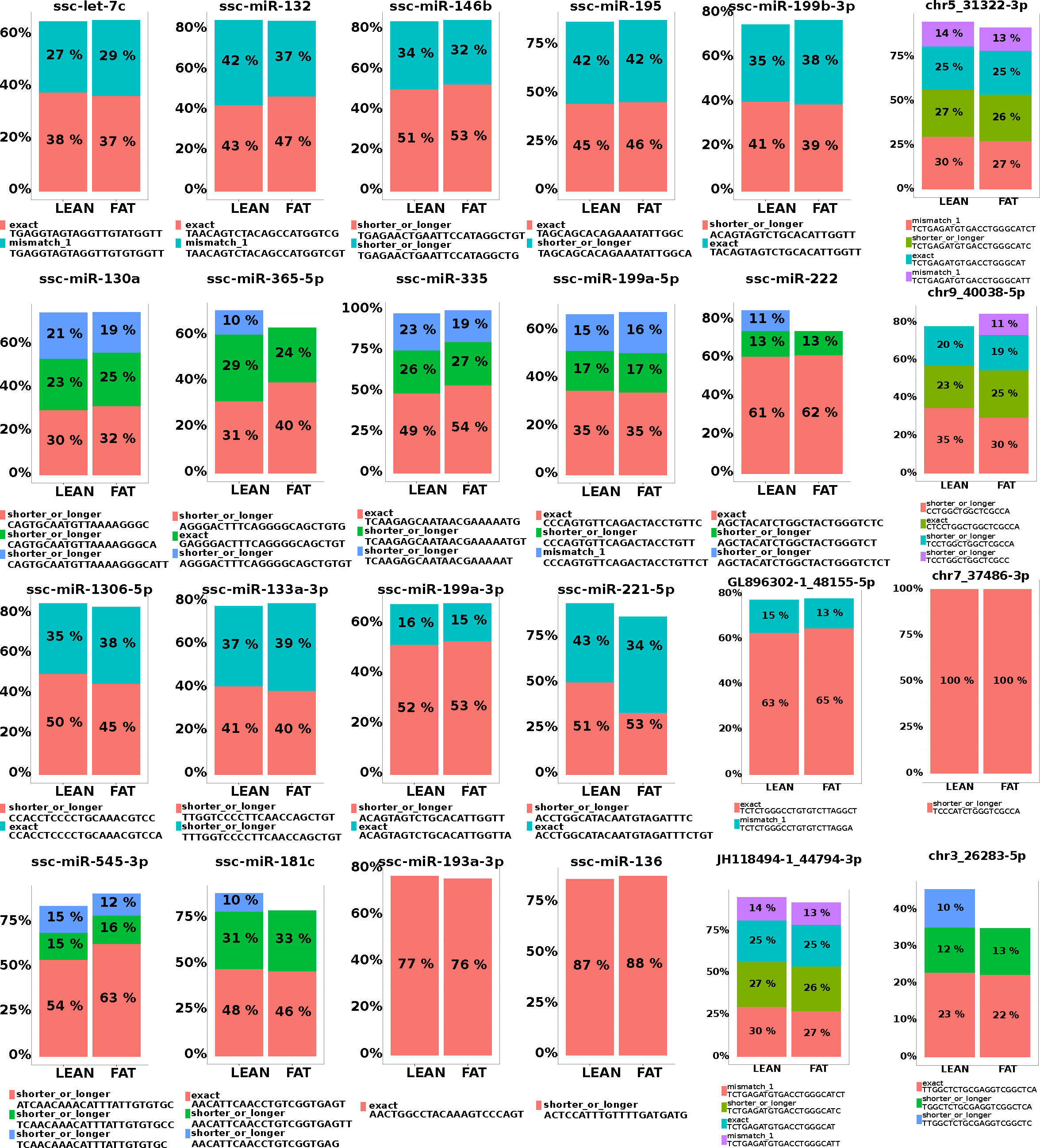
IsomiR composition of known and newly predicted miRNAs resulting differentially expressed in backfat tissue of LEAN and FAT animals. Colors and labels are relative to each single bar-chart. The same color may refer to different labels. Portion of the bars not reaching 100% represents isoforms composing each less than 10% of the overall miR expression. File: FigureSF3_rev.pdf

